# MULAN: Multimodal Protein Language Model for Sequence and Structure Encoding

**DOI:** 10.1101/2024.05.30.596565

**Authors:** Daria Frolova, Marina A. Pak, Anna Litvin, Ilya Sharov, Dmitry N. Ivankov, Ivan Oseledets

## Abstract

Most protein language models (PLMs), which are used to produce high-quality protein representations, use only protein sequences during training. However, the known protein structure is crucial in many protein property prediction tasks, so there is a growing interest in incorporating the knowledge about the protein structure into a PLM. In this study, we propose MULAN, a MULtimodal PLM for both sequence and ANgle-based structure encoding. MULAN has a pre-trained sequence encoder and an introduced Structure Adapter, which are then fused and trained together. According to the evaluation on 7 downstream tasks of various nature, both small and medium-sized MULAN models show consistent improvement in quality compared to both sequence-only ESM-2 and structure-aware SaProt. Importantly, our model offers a cheap increase in the structural awareness of the protein representations due to finetuning of existing PLMs instead of training from scratch. We perform a detailed analysis of the proposed model and demonstrate its awareness of the protein structure. The implementation, training data and model checkpoints are available at https://github.com/DFrolova/MULAN.

## 1 Introduction

Proteins, as unbranched heteropolymers, play a pivotal role in nearly all biological functions [1]. Comprising 20 distinct amino acids, the specific sequence of these amino acids determines the complex three-dimensional (3D) structure of the protein [2]. Subsequently, this 3D configuration governs the protein’s function [1]. Advances in genome sequencing initiated the growth of the number of publicly available protein data, unveiling a vast resource for understanding the molecular basis of life. The application of modern machine learning techniques for a better understanding of protein sequences can boost the development of diverse fields such as drug discovery, protein design, and biotechnology.

The abundance of protein sequences and their text-like nature made it possible to apply top-performing approaches from natural language processing to proteins. It is tempting to expect that protein sequence information alone would be sufficient for PLMs to infer protein structure and function. Recently, large protein language models (PLMs), such as ProtTrans [3], ESM-2 [4], and Ankh [5] have made remarkable progress in protein representation learning, surpassing previous approaches across various downstream tasks. However, it appears that the representation abilities of sequence-only PLMs are limited and some kind of structural information should be encoded directly into PLM. This limitation is represented by a significantly better performance [6] of structure-infused Alphafold [7] compared to sequence-only ESMFold [6].

At the same time, structural information about a huge amount of proteins has also become easily available with the revolutionary method AlphaFold [7]. Recently, several structural protein language models (SPLMs) were proposed. For example, SaProt [8] has successfully added some knowledge about the protein structure into the model, showing a better performance compared to sequence-only PLMs. However, existing SPLMs use only a small portion of structural information leaving room for improvement by extending the use of the protein 3D structure.

In this paper, we explore the potential of infusing existing PLMs with the protein structure information via small Structure Adapter. Our main contributions are:

- We introduce MULAN, a novel MULtimodal PLM for both sequence and ANgle-based structure processing. We propose the Structure Adapter, a MULAN module that uses residue torsion angles to represent the protein structure. Our model can work on top of existing PLMs and requires PLM finetuning, so it offers a cheap increase of structural awareness due to avoiding training from scratch.
- We evaluate the obtained structure-aware protein representations on a wide range of down-stream tasks. We show that MULAN performs significantly better compared to the corresponding sequence-only ESM-2 and shows improvement over structure-aware SaProt. For example, for the protein-protein interaction, our model shows improvement in AUROC by .055 for MULAN-small, by .031 for MULAN-ESM2, and by .048 for MULAN-SaProt.
- We perform an extensive ablation study to highlight the effectiveness and demonstrate the structural awareness of MULAN embeddings (see Section 4.2 and Figure 1).

## 2 Method

### 2.1 MULAN architecture

#### Structural information

In this study, we propose MULAN, which is a MULtimodal encoder PLM for both sequence and ANgle-based structure processing. MULAN uses the pre-trained base PLM and has the Structure Adapter – a module we introduce to incorporate the knowledge about the 3D protein structure (see Fig. 2). In our experiments, we use ESM-2 architecture, initializing the base PLM from ESM-2 or SaProt models. However, MULAN can be based on other PLMs.

**Figure 1:**
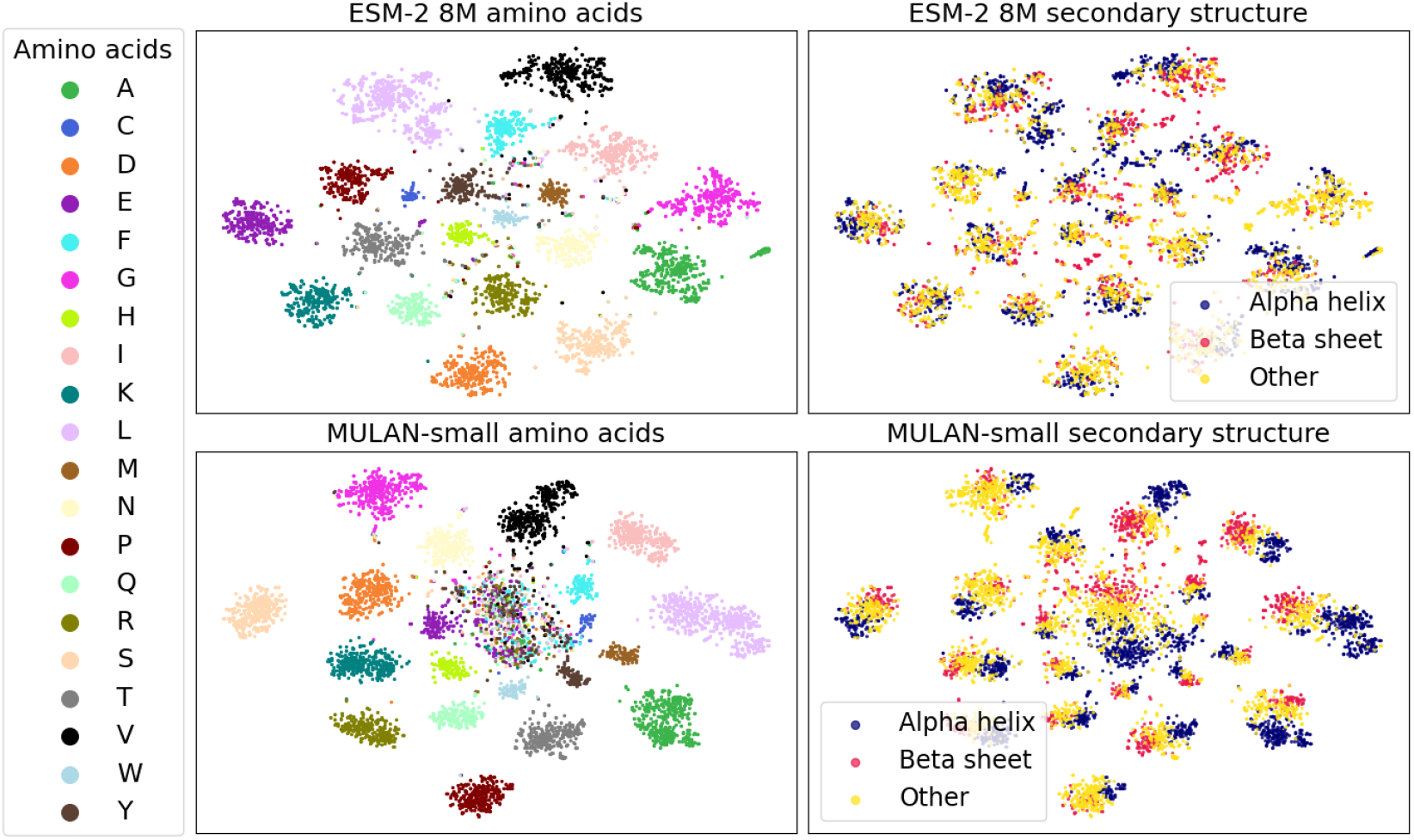
Visualization of residue embeddings of MULAN-small and ESM-2 8M on CASP12 dataset. We use different colors for amino acid residue types (left) and for the 3 states of secondary structure (right). The details of the experiment are presented in Section 4.3.

**Figure 2:**
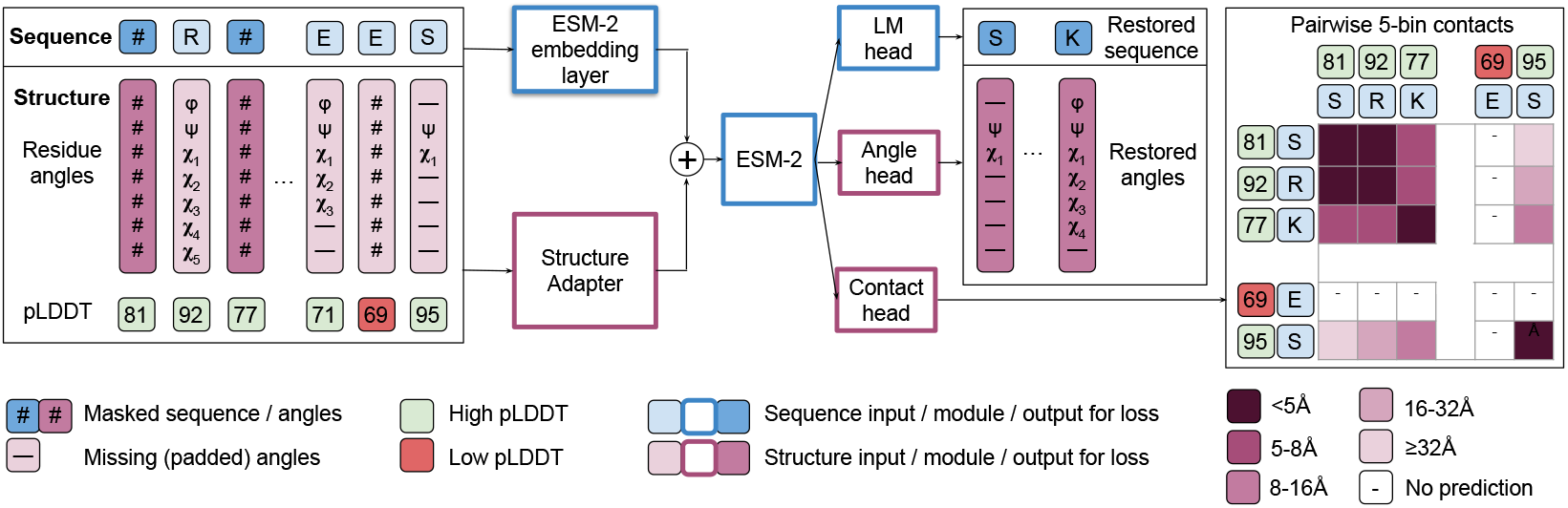
The proposed MULAN architecture. MULAN processes sequence inputs with the ESM-2 embeddings module, while structure inputs are passed to the Structure Adapter. Finally, both sequence and structure embeddings are summed up and passed to the ESM-2 model, which is then finetuned. Note that the Angle head and Contact head are optional. The base MULAN model utilizes only the LM head. Sequence-only ESM-2 modules are initialized from the pre-trained ESM-2 checkpoint and are shown in blue. Structure processing modules are shown in pink.

We use the information about the protein backbone torsion angles *ϕ* and *ψ* together with all residue side chain torsion angles *χ*_*i*_ (up to five *χ* angles for arginine). Missing *χ* angles together with undefined terminal *ϕ* and *ψ* angles are filled with the reserved value of 4 radians, which results in an angle vector [*ϕ, ψ, χ*_1_, *χ*_2_, *χ*_3_, *χ*_4_, *χ*_5_] for each residue. Moreover, we experiment with the contact prediction head, so we use pairwise distances between all residues in the form of a distance matrix as another source of structural data. Both residue angles and distance matrices are rotation- and translation-invariant; thus, they are easy to use inside a transformer model.

#### Structure Adapter

The introduced Structure Adapter is used to support the multimodality of our model and to fuse structural information with the sequence-only ESM-2 model. The Structure Adapter is a small encoder for the protein structure, which consists of an MLP followed by a Transformer layer. The MLP projects the residue angle vector to the residue angle embedding with dimension *h*, while the Transformer layer processes all protein angle embeddings at once (see Fig. 3). Given a protein of length *N*, the Structure Adapter returns an *N* × *h* angle embedding, or a structure bias. Finally, angle embeddings are added to initial ESM-2 residue embeddings of the same dimension *h* as a structure bias. The resulting structure-aware residue embeddings are then passed through the ESM-2 model. Due to the small size of the Structure Adapter, it does not add an overhead to the training or inference time compared to the base PLM.

**Figure 3:**
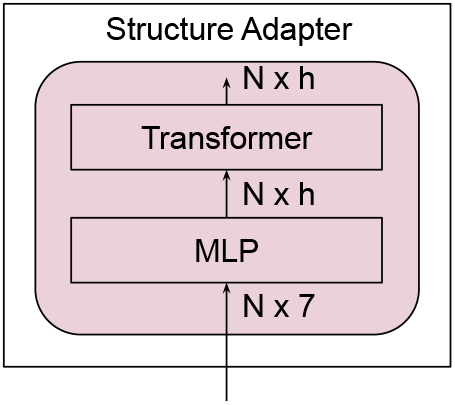
The architecture of the Structure Adapter.

#### Structure features prediction heads and objective function

One can use the knowledge about the protein structure not only as model input but also in the additional loss functions. We have experimented with two extra heads for the prediction of protein structure features, which were added to MULAN to increase its structural awareness. Firstly, we train to restore masked angle inputs similarly to the original sequence MLM objective. This is done using the Angle Prediction Head, which has the same architecture as the ESM-2 language modeling (LM) head except for the output dimension, which is 7, the number of residue angles. We use mean squared error (MSE) as a loss function.

Moreover, we use a residue-residue distance matrix inside the Contact Prediction Head. We binarize distances into 5 bins separated by the following distances: {5Å, 8Å, 16Å, 32Å}. Further, we predict *N* × *N* contact matrix based on the *N*_heads_ *\* N*_layers_ × *N* × *N* model attention weight tensor from all *N*_layers_ layers with *N*_heads_ attention heads. We use cross entropy (CE) loss and compute it only for residues with confident AlphaFold predictions to avoid training on noisy data. For each residue, AlphaFold produces an estimate of its confidence with a scale from 0 to 100 – predicted local distance difference test score (pLDDT). This measure corresponds to the AlphaFold predicted score on the lDDT-C*α* metric [9]. Predictions with pLDDT less than 70 are considered low confidence, so we do not compute contact loss on these residues. To sum up, we use the following objective function *ℒ* during training:

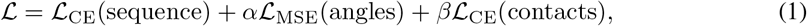

where *α* and *β* are floating point training hyperparameters. For base MULAN training, we do not use any extra heads, so *α* = *β* = 0. However, in the experiments that use structure features prediction heads, we utilize *α* = 5 and *β* = 0.5 as they have shown the best performance on downstream tasks. We discuss the impact of the addition of Contact and Angle Heads into MULAN in Section 4.

### 2.2 Training procedure and structure masking

We initialize the base model with a pre-trained ESM-2 checkpoint, so during training, we aim to finetune ESM-2 using additional structure inputs. We utilize the same masked language modelling objective [10] as ESM-2 for sequences, randomly masking 15% of tokens in a batch. In this work, we apply a similar masking strategy to the Structure Adapter. Residue angles are masked or replaced jointly with the corresponding residue letters. 80% of the time, angles are masked with a pre-defined mask value of −4 radians, which are then passed to the Structure Adapter. 10% of the time, the angle vector is replaced by another angle vector from the same protein, while in the rest cases, residue angles remain unchanged. Note, that both reserved angle values for padding and masking can have arbitrary values with absolute values higher than *π* in order not to mix with real angle values.

Moreover, we found out that only reliable structural information should be passed to the model. We observed that passing low-confidence predictions into the model worsens its performance (see Section 4.1). Therefore, we mask all residue angle vectors with AlphaFold pLDDT score lower than 70 (see Fig. 2).

## 3 Experiments

### 3.1 Training details

#### Training dataset

We use protein structures for Swiss-Prot proteins from AlphaFold Protein Structure Database (AlphaFold DB) [11] for the pre-training stage (503, 724 structures). We filter out proteins with a length of less than 30 amino acids. The final set comprises 501, 348 proteins. Following [12], we have randomly selected the validation set of 5000 proteins. Another training dataset was taken as is from ProstT5 setup [13]. It contains 17M AlphaFold protein structures not shorter than 30 residues. We refer to the datasets as AF-0.5M and AF-17M, respectively. Originally, all experiments with small models (8M parameters) were done using AF-0.5M, and the experiments from medium-sized models (35M parameters) were done using AF-17M.

#### Training procedure

The small MULAN model (referred to as MULAN-small) was initialized from the ESM-2 8M checkpoint and was trained on AF-0.5M dataset. For the medium-sized models, we use either ESM-2 35M or SaProt 35M checkpoint. We refer to these models as MULAN-ESM2 or MULAN-SaProt. The training of medium models was run on AF-17M dataset. During the pre-training stage, we randomly crop proteins that are longer than 1022 residues to the length of 1022. We follow [12] and use dynamic batch size without the concatenation of proteins along the sequence dimension during training. Also, to fully utilize GPU and minimize the amount of padding, we use sorted batching with dynamic batch size as in [14]: we keep a fixed number of tokens in the batch and form the batch from proteins with similar lengths. The extended training details and hyperparameters are detailed in Appendix A.

### 3.2 Downstream tasks

We tested our model on 8 downstream tasks (see Table 1), of which the first 7 are main tasks and the last one, Secondary structure prediction, served as a sanity check to verify the MULAN’s structure awareness. For the prediction of the protein Thermostability, Metal Ion Binding, Gene Ontology (GO), and Human Protein-Protein Interaction (HumanPPI) we follow the setup of SaProt [8]. Fluorescence prediction was taken from Ankh [5] setup. For the Secondary structure prediction task, we used TAPE [15] benchmark.

**Table 1:** Downstream tasks

| Task name | Prediction level | Task type | Evaluation metric | Data split sizes |
| --- | --- | --- | --- | --- |
|  |  |  |  | train / valid / test |
| Thermostability | protein | regression | SCC | 5, 056 / 639 / 1, 336 |
| Fluorescence | protein | regression | SCC | 21, 446 / 5, 362 / 27, 217 |
| Metal Ion Binding | protein | classification | AUC | 5, 066 / 662 / 665 |
| HumanPPI | protein pair | classification | AUC | 26, 317 / 234 / 180 |
| GO CC | protein | classification | $F_{\max}$ | 26, 225 / 2, 904 / 3, 350 |
| GO MF | protein | classification | $F_{\max}$ | 26, 225 / 2, 904 / 3, 350 |
| GO BP | protein | classification | $F_{\max}$ | 26, 225 / 2, 904 / 3, 350 |
| Secondary structure | residue | classification | accuracy | 8, 678 / 2, 170 / 21; 115; 434 |

AlphaFold structures for proteins from the datasets were retrieved from AlphaFold DB by Uniprot accession number if available. PDB IDs were mapped to UniProt accession numbers and retrieved from AlphaFold DB. If no protein identifier was available or there was no AlphaFold model for the UniProt accession number in the database, the protein was modeled by the standalone version of AlphaFold. For all described datasets we keep the original data splits. For the Secondary structure prediction, we use original experimental PDB structures for evaluation. The description of datasets and processing steps is detailed in Appendix B.1.

To evaluate all binary classification tasks we use the area under the ROC curve (AUC); for multiclass classification we measure accuracy; for multilabel GO annotation task, we follow [16] and use F_max_ score; and for the regression tasks we measure Spearman’s correlation coefficient (SCC).

### 3.3 Downstream task evaluation

#### Downstream model

To evaluate the performance of SPLMs, we extract protein embeddings from the last layer of a model. For protein-level tasks, we perform the average pooling of embeddings. Downstream task prediction is done using the model with the Light Attention architecture [17], which was designed to work with protein embeddings and shows better results than an MLP. We detail the architecture of the downstream model in Appendix B.2

#### Experimental setup

We train the downstream model and select optimal hyperparameters for each downstream task and embedding model independently based on the metric on the validation set. We set a fixed grid of hyperparameters for all downstream tasks, which includes the dropout rate and intermediate representation sizes of a downstream model. The grid and all used hyperparameters are detailed in Appendix B.3. We use AdamW optimizer (*β*_1_ = 0.9, *β*_2_ = 0.999) with a fixed learning rate (in most setups it is 5 *·* 10^−5^). The batch size is equal to 8192 for all experiments. The model is trained for 200 epochs, and we select the intermediate checkpoint with the best validation metric.

## 4 Results and discussion

### 4.1 Results

We train MULAN on top of three pre-trained models: ESM-2 8M, ESM-2 35M and SaProt AF 35M. We did not train MULAN for large models due to computational constraints; thus, we take only small and medium-sized PLMs for comparison. Our model is compared to ESM-2, the top-performing PLM, and SaProt, the best existing structural PLM. The quality of the protein representations on the considered downstream tasks is reported in Table 2. Note that we report results for small and medium-sized PLMs in separate parts of the table and highlight the best results in bold also separately according to the model size. Also, we do not report error bars for the results due to the computationally expensive process of re-training MULAN. The results for various large PLMs and SPLMs are presented in Appendix C. Our main findings are discussed below.

**Table 2:** Comparison of the performance of various PLMs and SPLMs on seven main downstream tasks. The first section corresponds to small models and the second – to medium-sized models. We indicate the best results in bold for each section separately

| Model name | Thermo- | Fluore- | Metal Ion | Human | GO |  |  |
| --- | --- | --- | --- | --- | --- | --- | --- |
|  | stability | scence | Binding | PPI | CC | MF | BP |
| | SCC ↑ | SCC ↑ | AUC ↑ | AUC ↑ | F<sub>max</sub> ↑ | F<sub>max</sub> ↑ | F<sub>max</sub> ↑ |
| **Small models** |  |  |  |  |  |  |  |
| ESM-2 8M | .666 | .579 | .731 | .698 | .490 | .529 | .400 |
| MULAN-small | **.672** | **.596** | **.778** | **.753** | **.492** | **.587** | **.426** |
| **Medium models** |  |  |  |  |  |  |  |
| ESM-2 35M | .689 | .592 | .793 | .751 | .489 | .621 | .443 |
| MULAN-ESM2 | .701 | .609 | .794 | **.782** | **.516** | **.636** | **.447** |
| Saprot AF 35M | .699 | .639 | .783 | .731 | .501 | .632 | .440 |
| MULAN-SaProt | **.704** | **.642** | **.800** | .779 | .505 | .631 | .442 |

#### MULAN outperforms sequence-only models

According to the results, MULAN offers a cheap improvement to the representation quality over all studied PLMs. The highest increase in performance is shown when MULAN is trained based on the sequence-only ESM-2. Both MULAN-small and medium-sized MULAN-ESM2 show consistent improvement over ESM-2 models of the corresponding size for all considered downstream datasets. The most significant boost is shown for HumanPPI (improvement by .055 in AUROC for MULAN-small and .031 for MULAN-ESM2) and molecular function prediction (improvement by .058 in F_max_ for MULAN-small and .015 for MULAN-ESM2). Also, fluorescence prediction quality increases by .017 in SCC for both MULAN models. This fact indicates that both protein property prediction and protein interaction prediction tasks benefit from the structural input. Moreover, the results show that MULAN is a general-purpose PLM that can produce high-quality results for protein downstream tasks of different nature.

#### MULAN further boosts SaProt

Even though SaProt already uses the protein structure information, MULAN-SaProt still increases the quality of protein representations on all but one considered downstream tasks. The best improvement of our model compared to SaProt is shown for HumanPPI and Metal Ion Binding tasks. At the same time, for GO MF the performance was kept at the original level of SaProt. These results highlight the fact that currently used Foldseek structural data are not enough to fully encode the protein structure. Still, there is room for improvement in the use of 3D structure, and the Structure Adapter has shown success in the further enrichment of protein representations with structural information.

#### MULAN offers cheap structural awareness

One more result is that MULAN-ESM2 performs better or comparable to SaProt in all considered downstream tasks except the Fluorescence prediction, where there is a significant dominance of SaProt. However, MULAN requires much less computations to achieve such results: we finetune the ESM-2 model for 1.9M steps with 32000 tokens per batch, which results in approximately 61B processed tokens during training. At the same time, SaProt requires training from scratch due to the complete change in the protein tokenization, which results in much longer training. SaProt reports that the computational cost of training is similar to ESM-1b [8], so for Saprot AF 35M we take the training time reported for ESM-2 35M. It was trained for 500K steps with 2M tokens per batch, which resulted in 1000B processed tokens in total. Overall, it is approximately 16 times faster to train MULAN compared to SaProt, which indicates that MULAN is a computationally cheap and effective method for inducing the protein structure information into the PLM.

### 4.2 Ablation study

In the experiments below we perform a detailed analysis of the performance of MULAN-small to reduce the amount of computations.

#### The Structure Adapter is the key factor for improvement

We show clear evidence of the importance of the Structure Adapter, not the structure features prediction heads alone (Contact and Angle) or the effect of our training dataset. We follow the same training procedure as with MULAN and perform three experiments. The results of the evaluation are presented in Table 3 in the first section with ESM-2. Firstly, we finetune pure ESM-2 8M on our training data (ESM-2 + finetune) to show that our training dataset and the finetuning procedure itself do not lead to a significant performance boost on downstream tasks. Secondly, we finetune ESM-2 with additional structure features prediction heads (ESM-2 + Angle / ESM-2 + Contact) without the Structure Adapter, showing that using the structure information in the loss function is not sufficient for improvement on downstream tasks: only metrics for HumanPPI increased significantly. Thus, we conclude that the Structure Adapter is the key ingredient in MULAN, which leads to an improvement in metrics for various downstream tasks. Moreover, experiments with the addition of Contact and Angle heads to MULAN do not show consistent improvement compared to MULAN alone (MULAN vs MULAN + Contact head vs MULAN + Angle head vs MULAN + Angle + Contact), so we decide that the base MULAN without any extra heads is the best setup, that unifies the simplicity of the approach with its effectiveness.

**Table 3:** Ablation study of the training pipeline for ESM-2 8M and MULAN-small. The first section corresponds to ESM-2 8M results without the Structure Adapter, while the second section – for MULAN-small. The best results for each section are shown in bold

| Model name | Thermo- | Fluore- | Metal Ion | Human | GO |  |  |
| --- | --- | --- | --- | --- | --- | --- | --- |
|  | stability | scence | Binding | PPI | CC | MF | BP |
| | SCC ↑ | SCC ↑ | AUC ↑ | AUC ↑ | F<sub>max</sub> ↑ | F<sub>max</sub> ↑ | F<sub>max</sub> ↑ |
| **ESM-2 8M** | .666 | **.579** | .731 | .698 | **.490** | .529 | .400 |
| + finetune | **.675** | .571 | **.747** | .706 | .488 | .537 | .409 |
| + Angle head | .660 | .569 | .704 | **.759** | .473 | **.544** | **.416** |
| + Contact head | .674 | .567 | .711 | .745 | .470 | .539 | .412 |
| **MULAN-small** | .672 | **.596** | .778 | .753 | **.492** | .587 | .426 |
| without pLDDT | .673 | .523 | .798 | .716 | .476 | .562 | .417 |
| + Angle head | .680 | .576 | .768 | .707 | .467 | .569 | .419 |
| + Contact head | **.690** | .567 | .745 | **.759** | .480 | **.594** | **.429** |
| + Angle + Contact | .675 | .532 | **.804** | .694 | .486 | .591 | .421 |
| sequence-only | .633 | .588 | .719 | .678 | .438 | .355 | .321 |
| from scratch | .651 | .475 | .687 | .725 | .436 | .458 | .349 |

#### Masking structural inputs with pLDDT for noise reduction

We show that it is useful to mask uncertain residues in AlphaFold structure models (with pLDDT *>* 70) before passing them to the Structure Adapter: see MULAN vs MULAN without pLDDT masking experiments(Table 3). Most of the downstream tasks benefit from this trick, most likely due to the noise reduction in input angles.

#### Starting from the pre-trained model is necessary

We demonstrate the benefits of finetuning the pre-trained ESM-2 model instead of training MULAN from scratch. Despite the addition of the structure bias to the initial ESM-2 embeddings, it is still possible and useful to adjust all model weights for new inputs and not to loose the base knowledge of a pre-trained model. We applied the same training procedure to the randomly initialized MULAN (see Table 3: MULAN vs MULAN from scratch), and the obtained results are much worse compared even to ESM-2 8M, which indicates the need for much more time for training MULAN from scratch.

The extended results with ablation studies and analysis of the used hyperparameters are shown in Appendix D.

### 4.3 Secondary structure prediction

We aim to show the awareness and proper use of 3D structure by MULAN. For this purpose, we evaluate our model on the secondary structure prediction downstream task (see Table 4). We report results on CASP12 [18], TS115 [19] and CB513 [20] datasets. Initially, MULAN was trained using AlphaFold structures. It masks uncertain residue predictions based on the AlphaFold pLDDT score. We evaluate the quality of secondary structure prediction using the initial datasets with experimental structures. For such type of structures, pLDDT is not available and applicable, so we pass angle information for all residues into MULAN without masking.

**Table 4:** Comparison of the performance of PLMs on the secondary structure prediction task. The table is split into sections based on the model size. The best results for each section are shown in bold

| Model name | 3-state, accuracy ↑ | | | 8-state, accuracy ↑ | | |
| --- | --- | --- | --- | --- | --- | --- |
|  | CASP12 | TS115 | CB513 | CASP12 | TS115 | CB513 |
| ESM-2 8M | .732 | .798 | .765 | .602 | .677 | .623 |
| MULAN-small | .889 | .909 | .887 | **.807** | **.842** | **.796** |
| MULAN-small + Contact head | **.893** | **.911** | **.888** | .802 | .838 | .793 |
| ESM-2 35M | .752 | .828 | .809 | .619 | .707 | .669 |
| MULAN-ESM2 | **.915** | **.932** | **.917** | **.839** | **.879** | **.844** |
| SaProt AF 35M | .900 | .924 | .910 | .805 | .848 | .817 |
| MULAN-SaProt | .905 | .927 | .912 | .807 | .852 | .818 |
| ESM-2 650M | .821 | .871 | .867 | .706 | .772 | .751 |
| ProstT5 1.2B | .858 | .887 | .891 | .747 | .800 | .797 |
| SaProt AF 650M | **.926** | **.948** | **.945** | **.844** | **.897** | **.869** |

#### Structural awareness of MULAN

According to the results from Table 4, all MULAN models demonstrate the awareness of the protein secondary structure. They surpass not only similar-sized ESM-2 models but also all existing large sequence-only PLMs by a large margin. Also, medium-sized MULAN models, especially MULAN-ESM2, show better structure awareness compared to SaProt, which indicates the usefulness of the proposed structure encoding using residue angles. Moreover, even a small MULAN utilizes the knowledge about the protein structure better than ProstT5 with 1.2B parameters. We do understand that the correct information about the secondary structure can be derived from angle inputs as well as from the Foldseek tokens used by SaProt. Hence, this experiment is done to demonstrate that MULAN actively uses the 3D structure.

#### Absence of 3D structure

One more check for the structural awareness of MULAN is to report its performance in the absence of the protein structure input. We show MULAN results with completely masked structural information in Table 3 (MULAN sequence-only experiment). The results show that input protein structure angles highly influence the quality of MULAN embeddings, resulting in even lower quality for a sequence-only MULAN compared to the initial ESM-2 8M model. This experiment shows the importance of structural information and its high usage by our model.

#### Visualization

The quality of the separation of the PLM representations according to some physical or structural property can serve as evidence of the model’s physical and structural awareness. We perform a comparison of MULAN-small and ESM-2 8M representations according to the visual quality of the t-SNE [21] visualization. We plot residue-level embeddings from the last layer of both models on the CASP12 dataset [18]. We highlight in color different amino acid residue types on the left and different secondary structure types (3 states) on the right (see Fig. 1). On the one hand, MULAN shows a much higher awareness of the secondary structure compared to ESM-2: for most amino acid clusters three secondary structure types are separated, while for ESM-2 they are mostly mixed up. On the other hand, MULAN does not lose the initial knowledge about the amino acid properties gained from the ESM-2 model. All these findings are in line with the pre-training strategy. ESM-2 has sequences as inputs, so it aligns amino acid representations in separate clusters. MULAN has both sequences and structure inputs; therefore, its representations are well-aligned in both domains.

## 5 Related work

### 5.1 Sequence-based models

Protein sequences are similar to human language: like letters are assembled into words that, in turn, form sentences, amino acids are chained into protein sequences that encode protein 3D structure which determines function. This resemblance makes it promising to apply best practices from the natural language processing field for solving protein-related tasks. Most PLMs are pre-trained with a masked language modeling (MLM) objective [10]: a part of the input sequence’s residues is randomly masked or replaced with other residues, and then the model aims to predict these corrupted tokens using the remaining sequence context. Models from the Transformer family (ESM-1b [12], ESM-2 [6], ProtTrans [3], Ankh [5]) have made huge progress in the protein representation learning. They show high performance in various downstream tasks, including but not limited to the prediction of protein secondary structure, residue contacts, sub-cellular localization, and the effect of mutation.

### 5.2 Structure-based models

The amino acid sequence solely defines the protein structure [2], which, in turn, defines all protein properties, including its function. However, the sequence alone is not sufficient enough for PLMs to infer all information about the protein [6]. Thus, attempts to infuse PLMs with structural context were made. To improve PLM’s capabilities, Zhang et al. [22] proposed to finetune ESM-2 on the remote homology detection task, which seems to implicitly incorporate protein structure-based features into the model. Indeed, the protein structure is more conserved than the protein sequence [23], and homology search being a more sensitive task than just finding similar protein sequences gives the model additional structural knowledge.

Recently, the idea to use protein 3D structure directly during the model pre-training has been given a lot of attention from the research community. ProstT5 [13], the first structure-augmented PLM, uses Foldseek, a special structural alphabet [24] (3Di) that describes the tertiary structure. As a result, each protein can be represented with either an amino acid sequence or a string of Foldseek letters of the same length that carries information about tertiary interactions. Incorporating that information into PLM was done by finetuning a sequence-only ProtT5 model to translate between the amino acid sequence and 3Di sequence to obtain structure-aware protein representations.

The graph-like tertiary structure of proteins gives ideas of infusing PLMs with structure information via Graph Neural Networks. In this setup, pre-training is left as MLM only [25, 26] or is augmented by Masked Structure Modeling task where not only parts of the sequence are masked but parts of the structure too [27, 28, 29]. In ESM-GearNet [30] it was proposed to fuse the protein sequence and structure information from state-of-the-art PLMs with graph structure encoders. The authors used various pre-training strategies including diffusion-based, and reported a performance boost compared to ESM-2 on several downstream tasks.

To the best of our knowledge, the most successful approach to using protein structure inside a PLM to date was shown in the SaProt [8]. Similarly to ProstT5, SaProt uses the 3Di alphabet to encode the structure. It represents each residue as a combination of amino acid and 3Di letters and is trained with MLM. This approach gives SaProt an improvement over ESM-2 on various downstream tasks.

## 6 Conclusion

In this paper, we propose MULAN, a novel multimodal 3D structure-aware protein language model for both sequence and structure processing. MULAN works on top of a sequence-only ESM-2 and has the introduced Structure Adapter that processes residue dihedral angles. Our model finetunes the pre-trained ESM-2 model offering a cheap incorporation of the knowledge about the protein structure into the model. We train small and medium-sized MULAN models and evaluate their protein representations on 7 downstream tasks. For all downstream tasks, our model demonstrates a consistent increase in performance compared to ESM-2 models, which were used for MULAN initialization. The best results were shown for the protein-protein interaction and the molecular function prediction. Moreover, our model demonstrates an improvement over SaProt when initialized from the SaProt model. Additionally, we indicate the importance of the proposed Structure Adapter, the main MULAN structure module. Finally, we demonstrate the structural awareness of our model in multiple experiments.

### Limitations and future work

Our model has one limitation: it was not trained on top of large PLMs because of the computational constraints. In the future work, we plan to train large MULAN. Additionally, we plan to investigate the effect of training on the experimental structures from the Protein Data Bank.

## A Training details

MULAN-small was trained with AdamW optimizer (*β*_1_ = 0.9, *β*_2_ = 0.999) [31] with learning rate linearly decaying from 10^−4^ to 2 *·* 10^−5^ during 100 epochs. We take 13000 tokens per batch for the base model and 3500 tokens per batch for a model with a contact head. The training process is run on 1 Tesla V100 GPU and lasts approximately 3 days (1.4M steps) for the base MULAN and 10 days (5.5M steps) for the setup with contact head.

For the medium-sized models were trained using 1 Tesla H100 GPU with 32000 tokens per batch with the use of AdamW optimizer (*β*_1_ = 0.9, *β*_2_ = 0.999). MULAN-ESM2 was trained for 15 epochs with the learning rate linearly increasing from 0 to 5 *·* 10^−5^ during the first 0.5 epochs and then decaying to 4 *·* 10^−5^ during the remaining training, which resulted in approximately 9 days (1.9M steps). MULAN-SaProt was trained for 10 epochs with the learning rate linearly increasing from 0 to 10^−5^ during the first 0.5 epochs and then decaying to 8 *·* 10^−6^ during the remaining training, which resulted in approximately 6 days (1.3M steps).

**Table 5:** Downstream task hyperparameters: learning rate (lr), dropout rate (dropout), intermediate representation sizes *h*_1_ and *h*_2_

| Task name | lr | dropout | $h_1$ | $h_2$ |
| --- | --- | --- | --- | --- |
| Metal Ion Binding | $5 \cdot 10^{-5}$ | {0.1, 0.2} | {1536, 1024, 512} | {512, 256} |
| HumanPPI | $5 \cdot 10^{-6}$ | 0.2 | {1536, 1024, 512} | {512, 256} |
| GO CC / MF / BP | $5 \cdot 10^{-4}$ | 0.1 | 1536 | 768 |
| Thermostability | $5 \cdot 10^{-5}$ | {0.1, 0.2} | {1536, 1024, 512} | {512, 256} |
| Fluorescence | $5 \cdot 10^{-5}$ | {0.1, 0.2} | {1536, 1024, 512} | {512, 256} |
| Secondary structure | $1 \cdot 10^{-4}$ | 0.1 | 1024 | 512 |

## B Downstream tasks

### B.1 Downstream datasets

We follow [8] and use their setup for protein Thermostability, Metal Ion Binding, GO, and HumanPPI. For Thermostability prediction, the “Human-cell” split from FLIP benchmark [32] is used. It relies on human data from Meltom atlas [33]. Also, one of the considered downstream tasks is Metal Ion Binding: we predict whether there are metal ion-binding sites in the protein [34]. The prediction of protein-protein interaction for human proteins (HumanPPI) [35] is taken from PEER benchmark [36]. We predict GO terms [16] and use all three branches independently: Molecular Function (MF), Biological Process (BP), and Cellular Component (CC). GO annotation is a multilabel prediction task. For all listed downstream tasks we use data provided by [8], so all used AlphaFold protein structures are available in the AlphaFold database.

Fluorescence prediction is done based on the data of the fluorescence intensity of green fluorescent protein (GFP) mutants [37]. We follow the setup of Ankh evaluation and use the split from TAPE [15] benchmark. We built an AlphaFold structure of the wild-type GFP protein and used Rosetta relaxation protocol [38] for the generation of mutant 3D structures. GFP_wt sequence was taken from the original dataset [37]. The reference GFP structure was provided to Rosetta to build mutant structures. Since these are single mutants, their structures should not differ a lot from the structure of wild-type GFP, and simple relaxation is enough. Then, we used pLDDT scores from the initial GFP structure for training on all mutant proteins.

Moreover, we evaluate our model on the secondary structure prediction task which is taken from TAPE benchmark [15]. We report results on three test datasets: CASP12 [18], TS115 [19] and CB513 [20], both for 3-state and 8-state setups. For this task, only experimental structures are available, so we use them as an input to MULAN. For all experimental structures, we pass all residue angles without masking into the MULAN. This is done because of the absence and inapplicability of pLDDT to the experimental structures.

### B.2 Downstream model architecture

Downstream task prediction is done using the model with the Light Attention architecture [17], which was designed to work with protein embeddings and shows better results than an MLP. The only difference is that we extend it by adding two extra intermediate layers *L*_1_ and *L*_2_: *L*_*i*_ = Dropout(ReLU(BatchNorm(Linear))) : 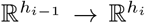, where *h*_1_ and *h*_2_ are the model hyperparameters, and *h*_0_ is the initial embedding dimension. They are added before the output Linear layer, which projects embeddings of size *h*_2_ into the downstream task target dimension.

### B.3 Downstream task hyperparameters

Here we present the grid used to select optimal hyperparameters for all downstream tasks (see 5). For the GO task, we have selected and fixed hyperparameters that perform well for all PLMs. We do not perform grid search because of the long time required for a single evaluation. Moreover, for GO it was optimal to increase the learning rate because of the much bigger output dimension in this task: up to 1943 classes for GO BP. For the HumanPPI task, the optimal learning rate differs from the base one due to the different nature of the task: we need to input two concatenated protein embeddings instead of one to the downstream model. Also, we reduced grid for HumanPPI (take 0.2 dropout rate) to decrease the number of required computations. The batch size is equal to 8192 for all experiments, and the training time is 200 epochs, but we select the intermediate checkpoint with the best validation metric. Since we use the secondary structure prediction task only to show the structural awareness of the model, we fix hyperparameters for the downstream task evaluation for a faster model evaluation.

## C Results for large models

We report the results of the evaluation of large PLMs and existing SPLMs in Table 6. The evaluation is performed similarly to previous sections using protein embeddings on all considered downstream tasks described in Appendix B.1.

**Table 6:** Comparison of the performance of various large PLMs and SPLMs on all downstream tasks. The best results are shown in bold

| Model name | Thermo- | Fluore- | Metal Ion | Human | GO |  |  |
| --- | --- | --- | --- | --- | --- | --- | --- |
|  | stability | scence | Binding | PPI | CC | MF | BP |
| | SCC ↑ | SCC ↑ | AUC ↑ | AUC ↑ | F<sub>max</sub> ↑ | F<sub>max</sub> ↑ | F<sub>max</sub> ↑ |
| ESM-2 650M | .694 | .601 | .781 | .754 | .523 | .678 | .479 |
| Ankh base 450M | .703 | .630 | **.837** | **.758** | .51 | .686 | .495 |
| Ankh large 1.2B | .668 | .638 | .787 | .738 | .517 | **.692** | **.501** |
| ProstT5 1.2B | .693 | .633 | .805 | .663 | .518 | .687 | .484 |
| Saprot AF 650M | **.711** | **.668** | .776 | .720 | **.540** | .658 | .464 |

## D Ablation studies

In this section, we present all ablation experiments that were performed during the selection of the training procedure of MULAN and hyperparameter tuning.

### D.1 Masking structural inputs with pLDDT

Table 7 shows experimental results with and without masking of structural inputs with low pLDDT score. Also, we report results with different pLDDT masking thresholds. The base one that we used was pLDDT *>* 70, a common threshold for the selection of good AF2 predictions suggested originally in AlphaFold2. Also, we compare it to masking with pLDDT *>* 50 as a filter of disordered regions. Overall, we conclude that for both MULAN and MULAN with contact head masking residue angle inputs with pLDDT *>* 70 is the best strategy.

**Table 7:** Comparison of the performance of MULAN with different pLDDT masking strategies. For each section of the table best results are shown in bold

| Model name | Thermo- | Fluore- | Metal Ion | Human | GO |  |  |
| --- | --- | --- | --- | --- | --- | --- | --- |
|  | stability | scence | Binding | PPI | CC | MF | BP |
| | SCC ↑ | SCC ↑ | AUC ↑ | AUC ↑ | F<sub>max</sub> ↑ | F<sub>max</sub> ↑ | F<sub>max</sub> ↑ |
| MULAN-small | **.673** | .523 | **.798** | .716 | .476 | .562 | .417 |
| + pLDDT 50 | .655 | .556 | .775 | .673 | .476 | .583 | .421 |
| + pLDDT 70 | .672 | **.596** | .778 | **.753** | **.492** | **.587** | **.426** |
| MULAN + Contact | .677 | .544 | **.780** | .696 | **.488** | .592 | **.432** |
| + pLDDT 70 + Cont. | **.690** | **.567** | .745 | **.759** | .480 | **.594** | .429 |

### D.2 Learning rate strategies

We used the same learning rate for all parameters of MULAN during training. However, we initialize the whole protein encoder from the ESM-2 pre-trained checkpoint, while the Structure Adapter is newly initialized. This fact suggests the possibility of using a smaller learning rate for ESM-2 modules compared to the Structure Adapter in order not to harm the pre-trained weights a lot. This idea was suggested and shown its effectiveness in ESM-GearNet paper [30], where they have a similar combination of randomly initialized and pre-trained modules. We follow the suggested setup and decrease the learning rate for ESM-2 modules by a factor of 10. According to the results, for MULAN there is no clear benefit of using the reduced learning rate for ESM-2 modules (see Table 8),so we decided to keep the learning rate constant for all MULAN modules to reduce the number of used hyperparameters.

**Table 8:** Comparison of the performance of MULAN-small with pLDDT masking with different learning rate (lr) strategies. If two learning rates are reported, the lowest corresponds to ESM-2 modules, while the highest – for the Structure Adapter. For each section of the table best results are shown in bold

| Model name | Thermo- | Fluore- | Metal Ion | Human | GO |  |  |
| --- | --- | --- | --- | --- | --- | --- | --- |
|  | stability | scence | Binding | PPI | CC | MF | BP |
| | SCC ↑ | SCC ↑ | AUC ↑ | AUC ↑ | F<sub>max</sub> ↑ | F<sub>max</sub> ↑ | F<sub>max</sub> ↑ |
| lr $5 \cdot 10^{-4} / 5 \cdot 10^{-5}$ | **.682** | .531 | .727 | **.754** | .483 | .559 | .414 |
| lr $1 \cdot 10^{-3} / 1 \cdot 10^{-4}$ | .674 | .557 | **.788** | .696 | .479 | .570 | .418 |
| lr $1 \cdot 10^{-4}$ for all | .672 | **.596** | .778 | .753 | **.492** | **.587** | **.426** |

### D.3 Number of layers in the Structure Adapter

We investigated the influence of the Structure Adapter size on the overall MULAN performance. The base version of MULAN has one Transformer layer, while one can easily increase their number. When choosing one layer for the Structure Adapter, we aimed to make it as small as possible to make it a lightweight additional component to ESM-2. However, we investigate the setup of a three-layer Structure Adapter for MULAN-small, which is based on ESM-2 8M. According to Table 9, there is no clear performance boost on the downstream tasks from the increase of the Structure Adapter size.

**Table 9:** Comparison of the performance of MULAN-small with the Structure Adapters of different sizes. For each section of the table best results are shown in bold

| Model name | Thermo- | Fluore- | Metal Ion | Human | GO |  |  |
| --- | --- | --- | --- | --- | --- | --- | --- |
|  | stability | scence | Binding | PPI | CC | MF | BP |
| | SCC ↑ | SCC ↑ | AUC ↑ | AUC ↑ | F<sub>max</sub> ↑ | F<sub>max</sub> ↑ | F<sub>max</sub> ↑ |
| MULAN 1-layer | .672 | **.596** | **.778** | .753 | **.492** | .587 | .426 |
| MULAN 3-layer | **.681** | .563 | .769 | **.763** | .486 | **.589** | **.427** |

### D.4 Additional experiments

#### Addition vs concatenation of embeddings

MULAN uses the structure bias from the Structure Adapter in a manner of positional embeddings: these structural embeddings are added to the main amino acid embeddings from the ESM-2 model. However, one may concatenate these embeddings instead of summing up. This approach leads to the different objectives used for amino acid embeddings (MLM) and structure embeddings (MLM for angle restoration). Also, concatenation leads to an increase in the length of the content passed to the Transformer model, causing significant memory and time overheads. The results of the experiment with the concatenation of embeddings did not show any benefit compared to the base setup.

#### Importance of angle masking

In our pre-training strategy, both amino acids and corresponding angle vectors are masked together. However, there are two other options that we have tested: independent masking of angles and residue letters and no angle masking at all. These experiments have shown worse results than the base approach. We explain by the fact that residue letters and their angle vectors are connected. For example, if the letter is masked, and the angle vector has only no side chain torsion angles defined, then the range of possible outcomes decreases from all 20 amino acids to only two: Glycine and Alanine. The opposite also holds: the known residue letter helps to restore the corresponding residue angle vector or at least the number of residue angles. Thus, we keep the joint masking strategy to force MULAN to learn as much information as possible.

